# Into the Void: Cavities and Tunnels are Essential for Functional Protein Design

**DOI:** 10.1101/2024.05.06.592825

**Authors:** Jiahui Zhang, Zhengxing Peng

## Abstract

The design of functional proteins is crucial as it enables the creation of tailored proteins with specific capabilities, unlocking the potential solutions to various biomedical and industrial challenges. The exact relationship between structure, sequence, and function in protein design is intricate, however, a consensus has been reached that the function of a protein is mostly decided by its structure, which further decides its sequence. While the integration of biology with artificial intelligence has propelled significant advancements in protein design and engineering, structure-based functional protein design, especially *de novo* design, the quest for satisfactory outcomes remains elusive. In this work, we use backbone geometry to represent the cavities and tunnels of functional proteins and show that they are essential for functional protein design. Correct cavity enables specific biophysical processes or biochemical reactions, while appropriate tunnels facilitate the transport of biomolecules or ions. We also provide a package called CAvity Investigation Navigator (CAIN) to help to do the analysis, which is available at https://github.com/JiahuiZhangNCSU/CAIN.

## 1 Introduction

Protein design is a technique by which proteins with desired properties are createdHuang et al. [2016], Pan and Kortemme [2021], Samish et al. [2011], which has been greatly and rapidly developed since the merge of biology and artificial intelligence (AI)Wang et al. [2018a], Notin et al. [2024]. Such a task becomes functional protein design if the desired property of the to-be-designed protein holds a specific biophysics or biochemistry function. Natural functional proteins involved over a significant period are bio-compounds that occur in every cellular function, exhibiting the remarkable efficiency in catalysis that drives the essential biophysics processes or biochemical reactions with exceptional speed and precisionMaynard Smith [1970], Walsh [2014], Agarwal [2006]. Despite the effectiveness of natural functional proteins like enzymes, protein design is pursued to enhance performance and create novel functionalities for various applications. Desired designed functional proteins offer optimized stability, specificity, efficiency, and immunogenicity, while also allowing for the development of tailored solutions to specific challengesGoldenzweig and Fleishman [2018], Kulp and Schief [2013]. Additionally, functional protein design enables the exploration of fundamental principles in biomolecular engineering, advancing our understanding of protein function and evolution. Protein function is decided by its structure, while protein structure is decided by its sequenceJohansson and Lindorff-Larsen [2018]. So, in most cases, the desired function and designed sequence are bridged by an optimized structure, thus the functional protein design breaks into two major steps: function design (from function to structure) and fold design (from structure to sequence)Listov et al. [2024]. In recent years, AI has significantly advanced protein designPaladino et al. [2017]. By analyzing vast protein datasets, AI algorithms have revolutionized protein design and engineering, enabling the creation of novel proteins with desired functions for diverse applicationsNotin et al. [2024]. However, despite these advancements, the achievements of AI in functional protein design are still not entirely satisfactory, especially in the field of *de novo* designListov et al. [2024].

In the field of fold design, machine learning algorithms such as ABACUS-RLiu et al. [2022a] and Protein-MPNNDauparas et al. [2022] have achieved remarkable accomplishments. However, such algorithms tend to give sequences that solely stabilize the structure and do not consider the protein function, so the successful case of *de novo* enzyme design based on such algorithms is rare. LigandMPNNKrishna et al. [2024] takes the protein binding the ligand into consideration in the task of fold design, which increases the performance of the design of ligand-binding proteins. On the other hand, machine learning-based fold design algorithms such as PSTChen et al. [2024] and CarbonDesignRen et al. [2023] integrate protein language modelsFerruz and Höcker [2022] into fold design, which dramatically improves the quality of designed sequences as the sequences naturally satisfy the protein grammars. For function design, machine learning algorithms such as RFdiffusionWatson et al. [2023] and SCUBA-DLiu et al. [2022b] are proven to be able to generate design-able backbone structures with given conditions (eg, a fixed pocket backbone geometry). RFdifussionAAKrishna et al. [2024] can further build protein backbone only based on a given ligand structure, which is a great step toward the binding protein and enzyme *de novo* design, especially for an artificial ligand or substrateKrishna et al. [2024]. Despite the methods being improved a lot since the age of AI, the inverse function problem is still limited in scopeListov et al. [2024]. Considering the general case of the data-driven protein design problem, as shown in equation1, we want the learned possibility function to be similar to the true possibility. Here *x* denotes the full information of a protein and the condition can be divided into two parts, the general condition (such as the stability and expressibility) and the specific condition (the desired biophysics or biochemical functions).

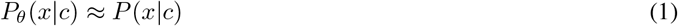

Suppose there is an ideal character vector (or generalized coordinates in physics) for a protein function characterized by data compression, the inverse function problem maximizes the learned possibility as shown in equation2, in which the *P*_*θ*_(*q*|*c*_*g*_, *c*_*s*_) part is of highly desired in a generation task (*c*_*g*_ is the general condition while *c*_*s*_ denotes the specific condition). From a physical perspective, there is a certain correlation between possibility and free energy, which states that 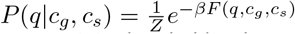, where *Z* is the partition function. If the system is in the equilibrium state, the free energy is solely decided by the stability (energy) and the protein space diversity (entropy) of a given *q*. This happens when all *c*_*s*_ are identical or we only consider the general condition in protein design, in other words, *P* (*q*|*c*_*g*_), or *F* (*q, c*_*g*_) = *F* ^*eq*^(*q*) is learnable by a large database with various and unbiased distributed *c*_*s*_. However, as the functions of each protein are greatly different, the system considered here is far from equilibrium. To characterize the difference of the possibility between true distribution and equilibrium distribution, we define 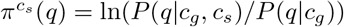. The physical significance of 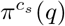 is straightforward as the ensemble average of it is exactly the total extropySivak and Crooks [2012], Gaveau et al. [2002] of the system: 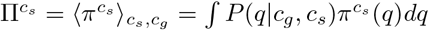, which is defined by the KL-divergence between the non-equilibrium distribution and the equilibrium distribution. The desired possibility can be written as equation3, from where we can see that the extropy is important in functional protein design.

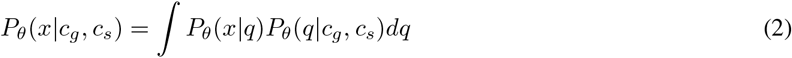

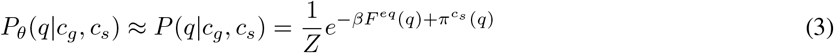

While the energy and entropy refer to the stability and protein space diversity of the given generalized coordinates, the extropy manifests the biofunction of the designed protein, such as the catalytic mechanism and transport pathway. If the extropy is not considered and the distribution is exactly Boltzmann’s distribution, then the designed proteins will be stable enough without desired bio functions, we refer to this phenomenon as “Boltzmann ruin”. In this work, we argue that focusing on the geometry of protein cavities is essential for understanding the catalytic mechanism and transport pathway and, thus is necessary for functional protein design. The study of protein cavities has a long historyLiang et al. [1998], Kuhn et al. [2006], Paramo et al. [2014]. However, the relationship between protein cavities and functional protein design, especially machine learning *de novo* functional protein design is still not fully disclosed. In this work, we discuss the importance of protein cavities such as ligand pockets and tunnels in protein functions, which may shed light on *de novo* functional protein design.

## 2 Methods

We use the alpha-spheresRooklin et al. [2015] defined by the Voronoi diagram of the protein atoms to represent protein cavities, which serve as a set of generalized coordinates of a protein. For *de novo* function design and fold design, we only consider the backbone *α*-Carbon atoms as we do not know the protein’s sequence, let alone the side chain conformation. Also, previous researches have shown that backbone conformation is important for protein pocket designLipsh-Sokolik et al. [2021], Lapidoth et al. [2018]. However, we encourage researchers to focus on backbone-based schedule for research other than *de novo* design as well, since in most cases, the side chains are much more flexible than the backboneTrbovic et al. [2009] and they can easily alter their conformation for adapting the ligand, which leads to a more rigid geometry for backbone-based cavities. An alpha-sphere can be defined by a 4D vector, in which the first three components describe the Cartesian coordinates of its center while the last component describes its radius. The pocket of a given ligand in a protein-ligand system is defined by a set of alpha-spheres whose center is within its radius to the nearest atom of the ligand. The non-overlapping volume of this set of alpha-spheres thereby defines the volume of the pocket. The ligand’s tunnel is defined as all the alpha-spheres that are closed enough (here we choose a cutoff of 2.8Å, which is the diameter of the water molecule) to the pocket alpha-spheres set. The length of the tunnel is defined by the furthest alpha-sphere to the geometry center of the ligand molecule along a pathway of centers of nearest alpha-spheres if the tunnel is a one-way tunnel while for the open tunnel, the length is defined by the largest distance along the centers of all the tunnel alpha-spheres. The alpha-spheres that are too large or too small are meaningless, for simplicity, we only consider alpha-spheres with a radius in the range of 3.0Å-6.0Å(for backbone-based schedule, the upper boundary might be higher, for example, 8Å).

We introduce inverse radius weighted distance (IRWD) to define the difference between two pockets. The definition of IRWD is the minimum of the square root of the inverse radius weighted sum of the nearest alpha-sphere distance of the two pockets, as stated in equation4, in which *ϵ* is the weight factor and *T*_*i*_ is any possible combination of translation and rotation operators. The larger the factor, the greater the effect of the radius is concerned. For those pockets with different numbers of alpha-spheres, we simply expend the smaller pocket to the same number of alpha-spheres as the larger pocket by appending pseudo-alpha-spheres at the center of the smaller pocket with an infinity radius. The exact value of the minimum is hard to obtain given two not-aligned pockets, we simplify the exact minimizing problem by first centering the two pockets according to their center Cartesian coordinates, then searching the rotation operator that gives the minimum IRWD by random sampling. Note that this is just one approach among many, and it might not be the optimal one. We look forward to future developments that could offer better solutions.

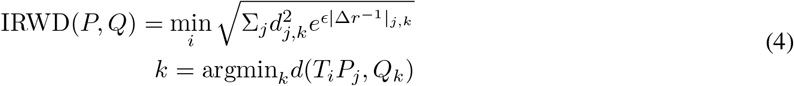

## 3 Results

Urate oxidaseWu et al. [1989] (Uox) (shown in figure1A), also known as uricase, is an enzyme found in some organisms, including certain bacteria, fungi, and mammalsOda et al. [2002]. It catalyzes the oxidation of uric acid to facilitate its excretion from the body. In humans and some primates, uricase activity is lacking due to a genetic mutationOda et al. [2002], leading to elevated levels of uric acid in the blood, which can contribute to conditions like goutVogt [2005]. As a result, uricase has been explored as a therapeutic agent for treating hyperuricemia and related disordersBomalaski and Clark [2004], Sherman et al. [2008]. Natural uricase exhibits high immunogenicity, rendering it unsuitable for direct therapeutic use in hyperuricemia-related disorders for humansCaliceti et al. [2001]. One possible solution is to reduce the immunogenicity by protein engineering based on natural uricaseNyborg et al. [2016], while *de novo* design is a more promising but more challenging way. To engineer or design uricase, researchers must comprehend its catalytic mechanism, which is closely related to the geometry of the cavity and tunnel of the target protein. The mechanism of uricase is shown in figure1B as proposed inGabison et al. [2008].

To investigate the geometry of the cavity and tunnel of uric acid in uricase, we employ the protocols described in section2 to visualize and analyze the cavity and tunnel, as shown in figure1C and figure1D. Most of the alpha-spheres of the pocket of uric acid in uricase overlap with the uric acid molecule, however, there are two notable clusters of alpha-spheres lying out of the uric acid molecule, which are labeled as red and yellow stars in figure1C. This is in agreement with the catalytic mechanism of uricase, in which the void space near the red star is required for the oxygen and water molecules that participate in the reaction while the void space near the yellow star is reserved for the water molecule that collaborates the reaction. Such void spaces are very important for the function design of enzymes since the correct size and shape of the protein cavities are the primary guarantee for a designed enzyme to have the desired function. The volume of uric acid pocket in the uric is calculated to be 899.14Å^3^ by CAIN. Based on the proper geometric properties of cavities, one can further design the physical and chemical properties of the cavities by fold design. In addition to cavities, the geometric properties of substrate tunnels also significantly influence enzyme functionality. These tunnels serve as pathways for substrates to reach the active site, dictating the kinetics of substrate binding and product releaseShan et al. [2022]. The geometry of these channels determines factors such as substrate diffusion rates, steric hindrance, and the orientation of substrates during catalysis. Optimal tunnel geometry facilitates efficient substrate binding and positioning, enhancing catalytic activity. Moreover, the precise arrangement of residues lining these tunnels can modulate substrate specificity and regulate enzyme activity. Thus, understanding and modeling the geometric properties of substrate tunnels are crucial for enzyme design and engineering. As shown in figure1D, the tunnel of substrate for uricase is a winding way that bridges the active site and the external environment. The length of the substrate tunnel for uricase is calculated to be 13.95Å by CAIN. Researchers may design artificial enzymes with desired kinetic properties by manipulating the tunnel length.

**Figure 1.**
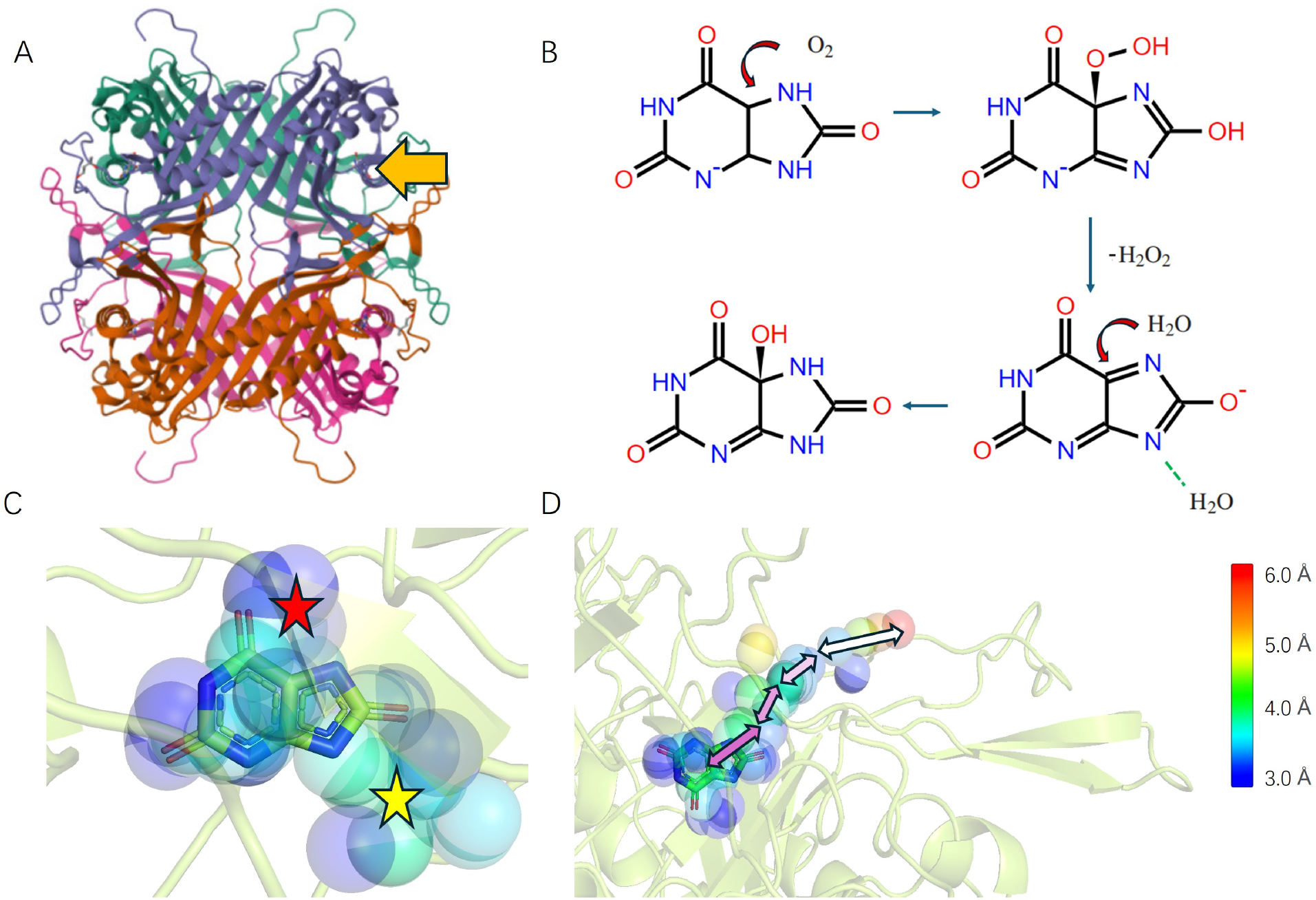
(A) The structure of Uox (PDB ID:4d12) with the active site noted by the arrow. (B) The mechanism of Uox catalytic reaction. (C) The cavity of the uric acid molecule in Uox and (D) the tunnel of the active site in Uox detected by CAIN, the color of the alpha-spheres represents the radius. The backbone of the protein is drawn by the transparent cartoon.

The other enzyme we focus on in this work is nucleoside 2-deoyribosyltransferase (NDT) (PDB ID: 1f8y, as shown in figure2A). This enzyme is vital in the salvage pathway of nucleotide metabolism in specific organisms. It facilitates the cleavage of *β*-2’-deoxyribonucleosides and then transfers the deoxyribosyl group to an acceptor purine or pyrimidine base and the proposed mechanismArmstrong et al. [1996] is given in figure2B. Studying such enzymes is valuable for elucidating cellular nucleotide metabolism, aiding drug development targeting these enzymes, and advancing biotechnological applications such as genetic engineeringArmstrong et al. [1996]. The rational or *de novo* design of enzymes with similar functions on artificial substrates is crucial for RNA vaccinesKramps and Elbers [2017], especially for the engineering of modified RNAHeidenreich et al. [1993]. Understanding how the mechanisms of natural nucleoside 2-deoyribosyltransferase constrain its structure is crucial for the design of artificial enzymes. One method involves focusing on protein cavities.

**Figure 2.**
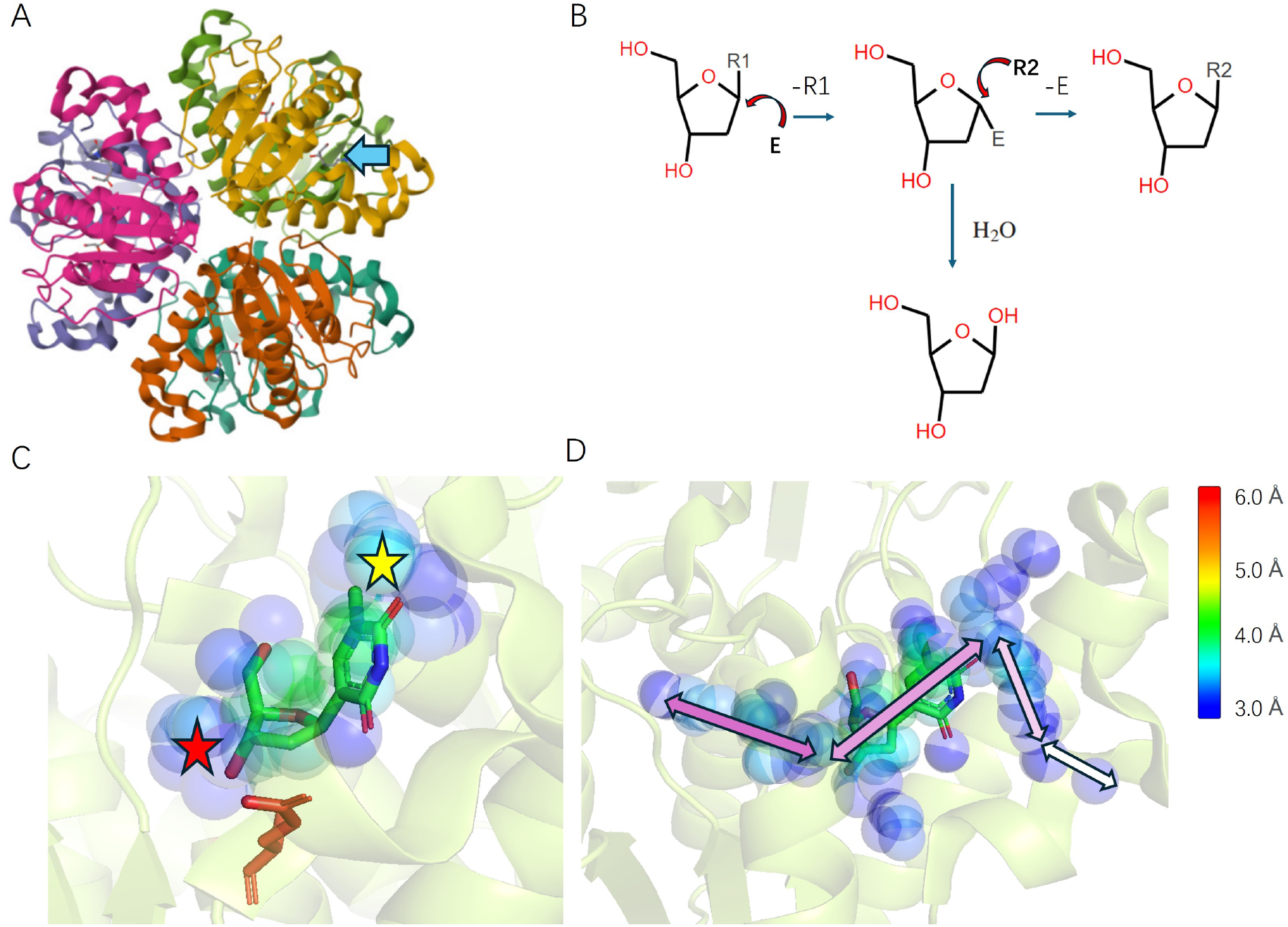
(A) The structure of NDT (PDB ID:1f8y) with the active site noted by the arrow. (B) The mechanism of NDT catalytic reaction. (C) The cavity of the nucleoside molecule in NDT and (D) the tunnel of the active site in Uox detected by CAIN, the color of the alpha-spheres represents the radius. The backbone of the protein is drawn by the transparent cartoon.

The pocket of the nucleoside molecule in NDT is extracted by CAIN, which is displayed in figure2C, in which the red stick labels the glutamine (E) that participates in the reaction. There are two main clusters of alpha-spheres outside the molecule, labeled by red and yellow stars. The void space near the yellow star is reserved for the purine since it requires more room than the pyrimidine, while the void space near the red star is prepared for a slightly left shifting of the substrate for the glutamine to reach the desired relative position to catalyze to reaction, which is demonstrated in previous molecular dynamics (MD) simulationsDel Arco et al. [2019]. The volume of this pocket is calculated to be 1200.49Å^3^ by CAIN. The more interesting part of NDT is its open zigzag tunnel with a length of 20.52Å calculated by CAIN, which is different from the one-way tunnel in Uox. The openness of the tunnel might be essential for the reaction, which should be one important constraint for *de novo* design of the artificial enzymes with similar functions on any modified nucleoside.

Our proposed protocol is not only valuable for enzymes but also for other functional proteins such as receptors. In this work, we investigate two neurotransmitter receptors, which are the dopamine D2 receptor and the *γ*-aminobutyric acid (GABA) A receptor. Dopamine and GABA are both important neurotransmitters in the nervous system. They have opposing effects, with dopamine mainly being responsible for excitatory functions (happiness) while GABA is primarily responsible for inhibitory functions (calmness)Iversen and Iversen [2007], Watanabe et al. [2002]. Understanding the receptors of dopamine and GABA is crucial as they play key roles in regulating various physiological and behavioral processes. Researching these receptors is particularly important in the fields of protein design and drug discovery. By elucidating the structure and function of dopamine D2 receptors and GABA_*A*_ receptors, scientists can gain insights into the mechanisms of neurotransmission and develop target therapies for neurological disorders such as Parkinson’s disease, schizophrenia, and bipolar disorderHsu et al. [2018]. Furthermore, knowledge of these receptors can aid in the design of novel drugs that modulate dopamine and GABA signaling pathways, offering potential treatments for a wide range of neurological and psychiatric conditionsBergman et al. [2000].

The structures we choose to analyze here are PDB ID:6cm4 (figure3A) and PDB ID: 6dw0(figure3C), in which the dopamine D2 receptor binds to an atypical antipsychotic drug molecule (risperidone) and the GABA_*A*_ receptor binds to the GABA moleculeWang et al. [2018b], Phulera et al. [2018]. The tunnels of both receptors are generated by CAIN as shown in figure3B and D. It can be seen that the tunnel of the dopamine D2 receptor is much deeper than the GABA_*A*_ receptor. This discrepancy holds significance as it affects the specificity and efficacy of ligand binding, thereby influencing downstream signaling cascades and neuronal activity modulation. Understanding these structural variances is crucial in pharmacology for both protein design and small molecule drug design. For instance, drugs targeting dopamine D2 receptors may aim for longer binding site tunnels to ensure optimal drug-receptor interactions, while those targeting GABA_*A*_ receptors may require shorter tunnels for effective modulation of proper neurotransmission. With a detailed description of tunnel geometry, elucidating the structural nuances of receptor-binding sites aids in the development of more precise and efficacious pharmacotherapies for various neurological and psychiatric disordersZheng et al. [2013], Sotriffer and Klebe [2002].

**Figure 3.**
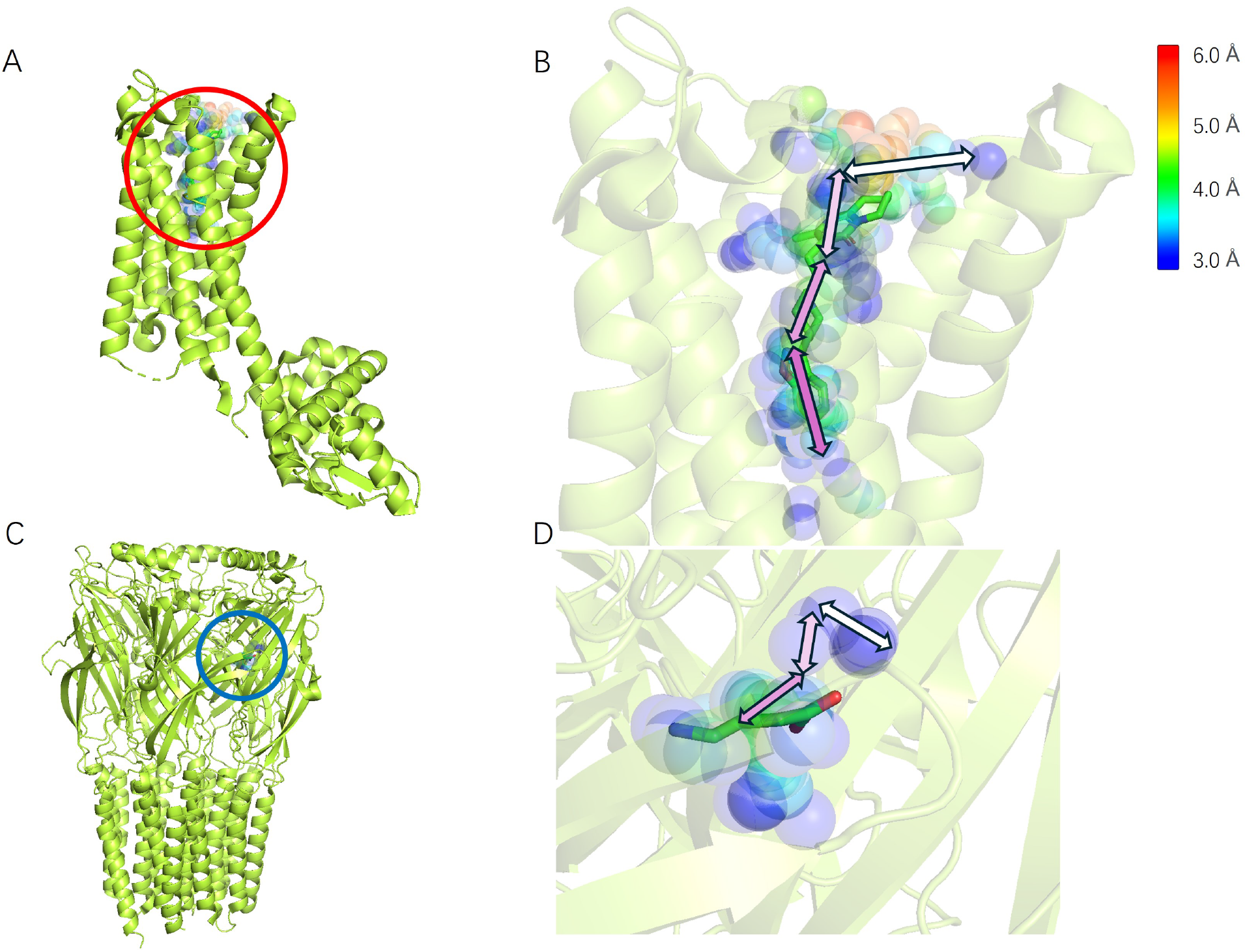
(A) The structure of the dopamine D2 receptor (PDB ID:6cm4) with binding site labeled in the red circle. (B) The zoomed-out view of the tunnel of the dopamine D2 receptor. (C) The structure of the GABA_*A*_ receptor (PDB ID:6dw0) with binding site labeled in the blue circle. (D) The zoomed-out view of the tunnel of the GABA_*A*_ receptor.

To further show the potential value of our protocol in functional protein design, we analyze the cavity and tunnel of a failure *de novo* design of artificial Uox. The design procedures follow the conventional steps, which are demonstrated in the left panel of figure4 A. To obtain the designed sequence with the desired function, the residues near the uric acid in the natural structure (PDB ID:4d12) is extracted, which is applied as the geometry constraint in the function design by SCUBA-DLiu et al. [2022b]. Then the generated backbone structure along with the fixed pocket residue type is utilized to obtain the designed protein sequence by ProteinMPNNDauparas et al. [2022]. To evaluate the quality of the designed sequence, AlphaFold2Jumper et al. [2021] is used to predict the structure of the designed protein, and SwissDockGrosdidier et al. [2011] is employed to do the molecular docking. Finally, we use CAIN to investigate the geometry of the cavity and tunnel of the designed protein.

The predicted structure with a binding uric acid molecule of the designed protein is shown in figure4C, which gives reasonable results in both structure prediction and molecular docking steps. However, this designed protein is found to have no ability to bind with uric acid, let alone being able to catalyze the desired reaction. One reason for this failure lies in the inappropriateness of its substrate cavity and tunnel. As shown in figure4B, the cavity of the designed protein and the cavity with the natural enzyme are significantly different, for some part of the uric acid, the designed protein even cannot offer void alpha-spheres with enough room, this is one possible reason that the designed protein cannot bind to the uric acid. Another possible reason is revealed by the analysis of tunnel geometry, which is shown in figure4C. The length of the tunnel is calculated to be 9.30Å, which is shorter than the natural tunnel, and the designed tunnel is much more twisting than the natural tunnel, which makes the uric acid extremely hard to go through and reach the pocket. The discussed failure in *de novo* design stems from a lack of consideration for the significance of the cavity and tunnel geometry in functional protein design.

**Figure 4.**
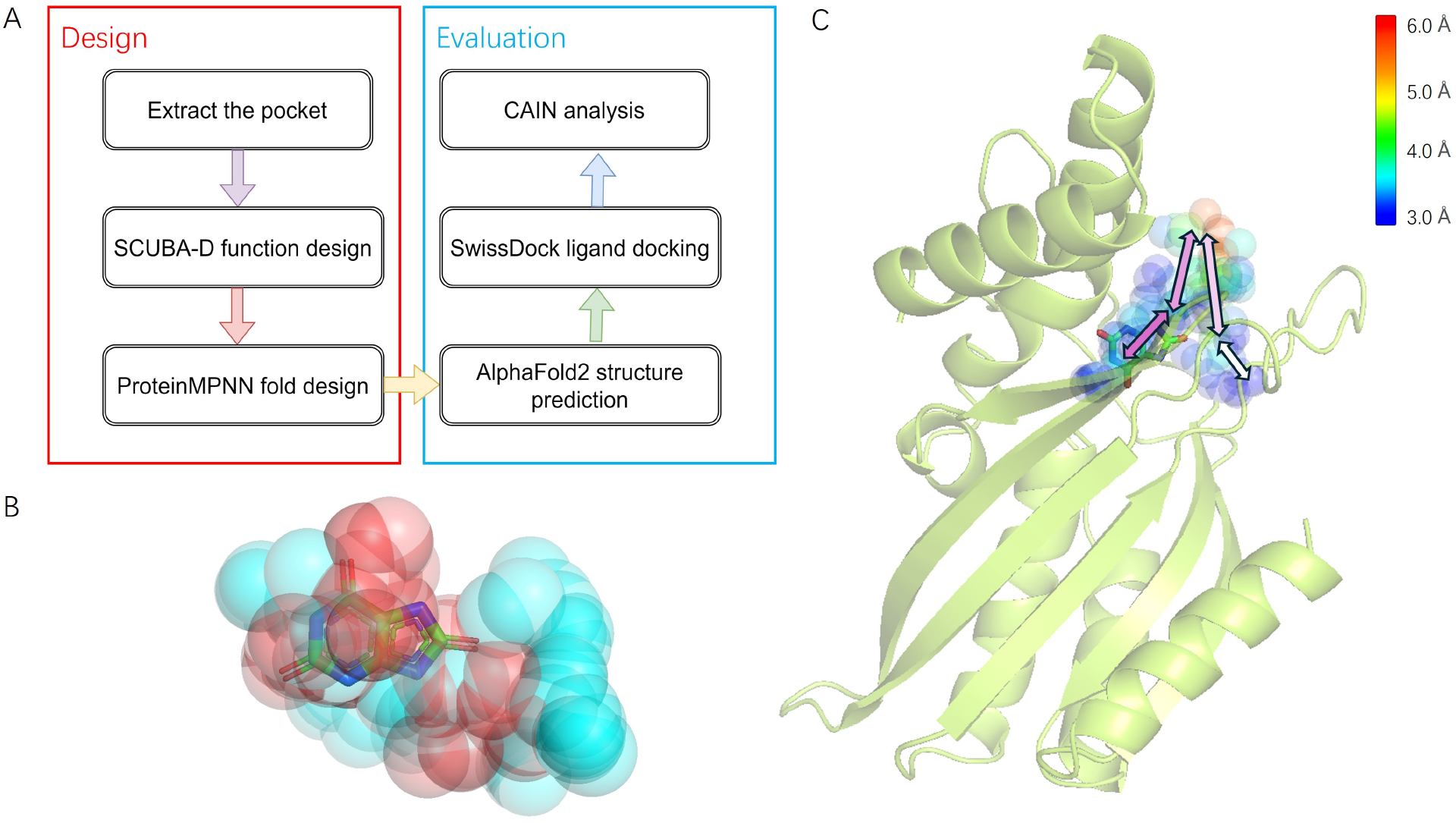
(A) The design and evaluation procedures of Uox design. (B) Comparison of the natural pocket (red) and the designed pocket (cyan). (C) The structure of the designed protein with the substrate tunnel is displayed.

## 4 Discussion

Our introduced methodology for analyzing the cavities and channels for functional proteins elucidates their significance in understanding the biochemical and biophysical functions of functional proteins such as enzymes and receptors. Our approach not only sheds light on the intricate mechanism underlying protein function but also provides crucial insights for the design of novel functional proteins. By delineating the structural features of protein pockets and pathways, we offer a comprehensive framework for rational protein engineering and drug design, thereby advancing our capabilities in harnessing protein functionality for various biomedical and biotechnological applications. The *de novo* design of functional protein is still a challengeDawson et al. [2019], Listov et al. [2024], Yang et al. [2021] due to the complicated relationship between protein structures and functionsSchulz and Schirmer [2013], Hegyi and Gerstein [1999]. Despite the significant achievements made in recent years through data-driven approachesChen et al. [2022], Liu et al. [2022a,b], Dauparas et al. [2022], Watson et al. [2023], the protein design using these methods still encounters issues where functionality is sacrificed for stability, leading to the Boltzmann ruin.

One way to overcome this challenge is to explicitly consider the cavities and tunnels in *de novo* protein design. In fold design, previous research shows that explicitly considering cavities in backbone generation and optimization is valuable for small molecule binding protein designXu et al. [2024], however, the choosing of the cavities is casual without an explicit mathematical definition. Furthermore, this approach fails to consider reaction mechanisms and substrate tunnels, thus limiting its application in enzyme design. By using the active site cavity detected by CAIN as a constraint, we anticipate improved performance in functional protein design. On the other hand, in graph neural networks (GNN) based fold design, it is possible to explicitly treat alpha-spheres of cavities and tunnels as nodes in the graph, with their coordinates and radius information. The IRWD defined in equation4 between the natural pocket and the designed pocket, together with the volume of the ligand cavity, can also be utilized as a score to rank the designed proteins.

This method of investigating cavities and tunnels may also apply to analyzing the MD trajectoriesSchmidtke et al. [2011]. The significance of cavity and tunnel dynamics for functional proteins lies in their crucial role in various biological processesBarbany et al. [2015], Paramo et al. [2014]. Cavities and tunnels within proteins serve as binding sites and pathways for ligands, substrates, and other molecules involved in biochemical reactions. The dynamics of these cavities and tunnels determine the accessibility and affinity of these binding sites, ultimately influencing the protein’s function. As the cavities and tunnels detected by CAIN are indeed sets of generalized coordinates of protein, one can expect that their motion obeys the generalized Langevin equationAyaz et al. [2021], Dalton et al. [2023] (GLE), which describes a non-Markovian process. However, in the dynamics of cavities and tunnels, there are occasions where it is necessary to introduce creation and annihilation operators, which adds complexity to the description of the dynamics. In any case, understanding the dynamics of cavities and tunnels in functional proteins is essential for elucidating their functional mechanisms, designing drugs that target specific binding sites, and engineering proteins with desired properties of various applications.

In conclusion, the method we propose for investigating protein cavities and tunnels offers clear mathematical definitions and straightforward implementation. We envision its utility in navigating the design of functional proteins, especially for the mechanism-based design. We also anticipate its positive impact on other structural biology-related fields. The code is available at https://github.com/JiahuiZhangNCSU/CAIN.

## Acknowledgement

We would like to thank Mingze Yang for sharing the sequence of the *de novo* designed uricase and its design strategy. We would like to express our sincere gratitude to Dr. Mengkai Feng for his invaluable guidance and insightful explanations of non-equilibrium statistical mechanics.

## References

Po-Ssu Huang, Scott E Boyken, and David Baker. The coming of age of de novo protein design. Nature, 537(7620): 320–327, 2016.

Xingjie Pan and Tanja Kortemme. Recent advances in de novo protein design: Principles, methods, and applications. Journal of Biological Chemistry, 296, 2021.

Ilan Samish, Christopher M MacDermaid, Jose Manuel Perez-Aguilar, and Jeffery G Saven. Theoretical and computational protein design. Annual review of physical chemistry, 62:129–149, 2011.

Jingxue Wang, Huali Cao, John ZH Zhang, and Yifei Qi. Computational protein design with deep learning neural networks. Scientific reports, 8(1):1–9, 2018a.

Pascal Notin, Nathan Rollins, Yarin Gal, Chris Sander, and Debora Marks. Machine learning for functional protein design. Nature Biotechnology, 42(2):216–228, 2024.

John Maynard Smith. Natural selection and the concept of a protein space. Nature, 225(5232):563–564, 1970.

Gary Walsh. Proteins: biochemistry and biotechnology. John Wiley & Sons, 2014.

Pratul K Agarwal. Enzymes: An integrated view of structure, dynamics and function. Microbial cell factories, 5:1–12, 2006.

Adi Goldenzweig and Sarel J Fleishman. Principles of protein stability and their application in computational design. Annual review of biochemistry, 87:105–129, 2018.

Daniel W Kulp and William R Schief. Advances in structure-based vaccine design. Current opinion in virology, 3(3): 322–331, 2013.

Kristoffer E Johansson and Kresten Lindorff-Larsen. Structural heterogeneity and dynamics in protein evolution and design. Current Opinion in Structural Biology, 48:157–163, 2018.

Dina Listov, Casper A Goverde, Bruno E Correia, and Sarel Jacob Fleishman. Opportunities and challenges in design and optimization of protein function. Nature Reviews Molecular Cell Biology, pages 1–15, 2024.

Antonella Paladino, Filippo Marchetti, Silvia Rinaldi, and Giorgio Colombo. Protein design: from computer models to artificial intelligence. Wiley Interdisciplinary Reviews: Computational Molecular Science, 7(5):e1318, 2017.

Yufeng Liu, Lu Zhang, Weilun Wang, Min Zhu, Chenchen Wang, Fudong Li, Jiahai Zhang, Houqiang Li, Quan Chen, and Haiyan Liu. Rotamer-free protein sequence design based on deep learning and self-consistency. Nature Computational Science, 2(7):451–462, 2022a.

Justas Dauparas, Ivan Anishchenko, Nathaniel Bennett, Hua Bai, Robert J Ragotte, Lukas F Milles, Basile IM Wicky, Alexis Courbet, Rob J de Haas, Neville Bethel, et al. Robust deep learning–based protein sequence design using proteinmpnn. Science, 378(6615):49–56, 2022.

Rohith Krishna, Jue Wang, Woody Ahern, Pascal Sturmfels, Preetham Venkatesh, Indrek Kalvet, Gyu Rie Lee, Felix S Morey-Burrows, Ivan Anishchenko, Ian R Humphreys, et al. Generalized biomolecular modeling and design with rosettafold all-atom. Science, page eadl2528, 2024.

Dexiong Chen, Philip Hartout, Paolo Pellizzoni, Carlos Oliver, and Karsten Borgwardt. Endowing protein language models with structural knowledge. arXiv preprint arXiv:2401.14819, 2024.

Milong Ren, Chungong Yu, Dongbo Bu, and Haicang Zhang. Highly accurate and robust protein sequence design with carbondesign. bioRxiv, pages 2023–08, 2023.

Noelia Ferruz and Birte Höcker. Controllable protein design with language models. Nature Machine Intelligence, 4(6): 521–532, 2022.

Joseph L Watson, David Juergens, Nathaniel R Bennett, Brian L Trippe, Jason Yim, Helen E Eisenach, Woody Ahern, Andrew J Borst, Robert J Ragotte, Lukas F Milles, et al. De novo design of protein structure and function with rfdiffusion. Nature, 620(7976):1089–1100, 2023.

Yufeng Liu, Linghui Chen, and Haiyan Liu. De novo protein backbone generation based on diffusion with structured priors and adversarial training. bioRxiv, pages 2022–12, 2022b.

David A Sivak and Gavin E Crooks. Near-equilibrium measurements of nonequilibrium free energy. Physical Review Letters, 108(15):150601, 2012.

B Gaveau, K Martinás, M Moreau, and J Tóth. Entropy, extropy and information potential in stochastic systems far from equilibrium. Physica A: Statistical Mechanics and its Applications, 305(3-4):445–466, 2002.

Jie Liang, Clare Woodward, and Herbert Edelsbrunner. Anatomy of protein pockets and cavities: measurement of binding site geometry and implications for ligand design. Protein science, 7(9):1884–1897, 1998.

Daniel Kuhn, Nils Weskamp, Stefan Schmitt, Eyke Hüllermeier, and Gerhard Klebe. From the similarity analysis of protein cavities to the functional classification of protein families using cavbase. Journal of molecular biology, 359 (4):1023–1044, 2006.

Teresa Paramo, Alexandra East, Diana Garzón, Martin B Ulmschneider, and Peter J Bond. Efficient characterization of protein cavities within molecular simulation trajectories: trj_cavity. Journal of chemical theory and computation, 10 (5):2151–2164, 2014.

David Rooklin, Cheng Wang, Joseph Katigbak, Paramjit S Arora, and Yingkai Zhang. Alphaspace: fragment-centric topographical mapping to target protein–protein interaction interfaces. Journal of chemical information and modeling, 55(8):1585–1599, 2015.

Rosalie Lipsh-Sokolik, Dina Listov, and Sarel J Fleishman. The abdesign computational pipeline for modular backbone assembly and design of binders and enzymes. Protein Science, 30(1):151–159, 2021.

Gideon Lapidoth, Olga Khersonsky, Rosalie Lipsh, Orly Dym, Shira Albeck, Shelly Rogotner, and Sarel J Fleishman. Highly active enzymes by automated combinatorial backbone assembly and sequence design. Nature communications, 9(1):2780, 2018.

Nikola Trbovic, Jae-Hyun Cho, Robert Abel, Richard A Friesner, Mark Rance, and Arthur G Palmer III. Protein side-chain dynamics and residual conformational entropy. Journal of the American Chemical Society, 131(2): 615–622, 2009.

XW Wu, Cheng Chi Lee, Donna M Muzny, and C Thomas Caskey. Urate oxidase: primary structure and evolutionary implications. Proceedings of the National Academy of Sciences, 86(23):9412–9416, 1989.

Masako Oda, Yoko Satta, Osamu Takenaka, and Naoyuki Takahata. Loss of urate oxidase activity in hominoids and its evolutionary implications. Molecular biology and evolution, 19(5):640–653, 2002.

Bruno Vogt. Urate oxidase (rasburicase) for treatment of severe tophaceous gout. Nephrology Dialysis Transplantation, 20(2):431–433, 2005.

John S Bomalaski and Mike A Clark. Serum uric acid-lowering therapies: where are we heading in management of hyperuricemia and the potential role of uricase. Current rheumatology reports, 6(3):240–247, 2004.

Merry R Sherman, Mark GP Saifer, and Fernando Perez-Ruiz. Peg-uricase in the management of treatment-resistant gout and hyperuricemia. Advanced drug delivery reviews, 60(1):59–68, 2008.

Paolo Caliceti, Oddone Schiavon, and Francesco M Veronese. Immunological properties of uricase conjugated to neutral soluble polymers. Bioconjugate chemistry, 12(4):515–522, 2001.

Andrew C Nyborg, Chris Ward, Anna Zacco, Benoy Chacko, Luba Grinberg, James C Geoghegan, Ryan Bean, Michaela Wendeler, Frank Bartnik, Ellen O’Connor, et al. A therapeutic uricase with reduced immunogenicity risk and improved development properties. PLoS One, 11(12):e0167935, 2016.

Laure Gabison, Thierry Prangé, Nathalie Colloc’h, Mohamed El Hajji, Bertrand Castro, and Mohamed Chiadmi. Structural analysis of urate oxidase in complex with its natural substrate inhibited by cyanide: mechanistic implications. BMC structural biology, 8:1–8, 2008.

Yibing Shan, Venkatesh P Mysore, Abba E Leffler, Eric T Kim, Shiori Sagawa, and David E Shaw. How does a small molecule bind at a cryptic binding site? PLoS computational biology, 18(3):e1009817, 2022.

ShellyR Armstrong, William J Cook, Steven A Short, and Steven E Ealick. Crystal structures of nucleoside 2-deoxyribosyltransferase in native and ligand-bound forms reveal architecture of the active site. Structure, 4(1): 97–107, 1996.

Thomas Kramps and Knut Elbers. Introduction to rna vaccines. RNA Vaccines: Methods and Protocols, pages 1–11, 2017.

Olaf Heidenreich, Wolfgang Pieken, and Fritz Eckstein. Chemically modified rna: approaches and applications. The FASEB journal, 7(1):90–96, 1993.

Jon Del Arco, Almudena Perona, Leticia González, Jesús Fernández-Lucas, Federico Gago, and Pedro A Sánchez-Murcia. Reaction mechanism of nucleoside 2-deoxyribosyltransferases: free-energy landscape supports an oxocarbenium ion as the reaction intermediate. Organic & Biomolecular Chemistry, 17(34):7891–7899, 2019.

Susan D Iversen and Leslie L Iversen. Dopamine: 50 years in perspective. Trends in neurosciences, 30(5):188–193, 2007.

Masahito Watanabe, Kentaro Maemura, Kiyoto Kanbara, Takumi Tamayama, and Hana Hayasaki. Gaba and gaba receptors in the central nervous system and other organs. International review of cytology, 213:1–47, 2002.

Wen-Yu Hsu, Hsien-Yuan Lane, and Chieh-Hsin Lin. Medications used for cognitive enhancement in patients with schizophrenia, bipolar disorder, alzheimer’s disease, and parkinson’s disease. Frontiers in psychiatry, 9:328047, 2018.

J Bergman, CP France, SG Holtzman, JL Katz, W Koek, and DN Stephens. Agonist efficacy, drug dependence, and medications development: preclinical evaluation of opioid, dopaminergic, and gaba a-ergic ligands. Psychopharmacology, 153:67–84, 2000.

Sheng Wang, Tao Che, Anat Levit, Brian K Shoichet, Daniel Wacker, and Bryan L Roth. Structure of the d2 dopamine receptor bound to the atypical antipsychotic drug risperidone. Nature, 555(7695):269–273, 2018b.

Swastik Phulera, Hongtao Zhu, Jie Yu, Derek P Claxton, Nate Yoder, Craig Yoshioka, and Eric Gouaux. Cryo-em structure of the benzodiazepine-sensitive α1β1γ2s tri-heteromeric gabaa receptor in complex with gaba. Elife, 7: e39383, 2018.

Xiliang Zheng, LinFeng Gan, Erkang Wang, and Jin Wang. Pocket-based drug design: exploring pocket space. The AAPS journal, 15:228–241, 2013.

Christoph Sotriffer and Gerhard Klebe. Identification and mapping of small-molecule binding sites in proteins: computational tools for structure-based drug design. Il Farmaco, 57(3):243–251, 2002.

John Jumper, Richard Evans, Alexander Pritzel, Tim Green, Michael Figurnov, Olaf Ronneberger, Kathryn Tunyasuvunakool, Russ Bates, Augustin Žídek, Anna Potapenko, et al. Highly accurate protein structure prediction with alphafold. Nature, 596(7873):583–589, 2021.

Aurelien Grosdidier, Vincent Zoete, and Olivier Michielin. Swissdock, a protein-small molecule docking web service based on eadock dss. Nucleic acids research, 39(suppl_2):W270–W277, 2011.

William M Dawson, Guto G Rhys, and Derek N Woolfson. Towards functional de novo designed proteins. Current Opinion in Chemical Biology, 52:102–111, 2019.

Che Yang, Fabian Sesterhenn, Jaume Bonet, Eva A van Aalen, Leo Scheller, Luciano A Abriata, Johannes T Cramer, Xiaolin Wen, Stéphane Rosset, Sandrine Georgeon, et al. Bottom-up de novo design of functional proteins with complex structural features. Nature Chemical Biology, 17(4):492–500, 2021.

Georg E Schulz and R Heiner Schirmer. Principles of protein structure. Springer Science & Business Media, 2013.

Hedi Hegyi and Mark Gerstein. The relationship between protein structure and function: a comprehensive survey with application to the yeast genome. Journal of molecular biology, 288(1):147–164, 1999.

Yaoxi Chen, Quan Chen, and Haiyan Liu. Depact and pacmatch: A workflow of designing de novo protein pockets to bind small molecules. Journal of Chemical Information and Modeling, 62(4):971–985, 2022.

Yang Xu, Xiuhong Hu, Chenchen Wang, Yongrui Liu, Quan Chen, and Haiyan Liu. De novo design of cavity-containing proteins with a backbone-centered neural network energy function. Structure, 2024.

Peter Schmidtke, Axel Bidon-Chanal, F Javier Luque, and Xavier Barril. Mdpocket: open-source cavity detection and characterization on molecular dynamics trajectories. Bioinformatics, 27(23):3276–3285, 2011.

Montserrat Barbany, Tim Meyer, Adam Hospital, Ignacio Faustino, Marco D’Abramo, Jordi Morata, Modesto Orozco, and Xavier de la Cruz. Molecular dynamics study of naturally existing cavity couplings in proteins. PloS one, 10(3): e0119978, 2015.

Cihan Ayaz, Lucas Tepper, Florian N Brünig, Julian Kappler, Jan O Daldrop, and Roland R Netz. Non-markovian modeling of protein folding. Proceedings of the National Academy of Sciences, 118(31):e2023856118, 2021.

Benjamin A Dalton, Cihan Ayaz, Henrik Kiefer, Anton Klimek, Lucas Tepper, and Roland R Netz. Fast protein folding is governed by memory-dependent friction. Proceedings of the National Academy of Sciences, 120(31):e2220068120, 2023.

